## Supplemental Information for "Regional structural-functional connectome coupling is heritable and associated with age, sex and cognitive scores in adults"

### Supplementary material

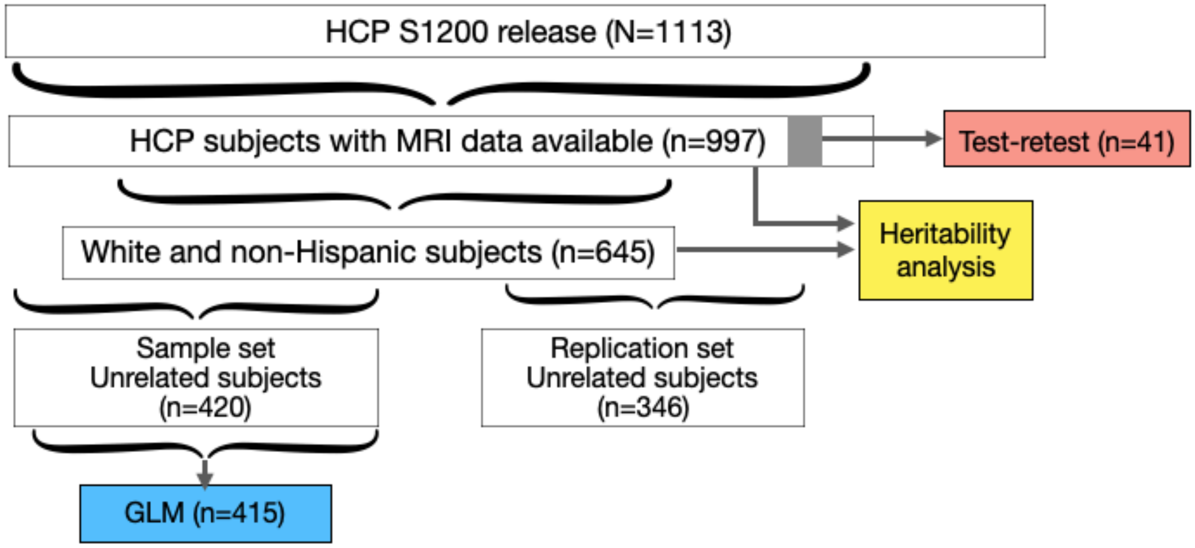


**Figure S1: Flowchart illustrating the selection of the HCP data for each of the analyses.** We began with the final S1200 HCP release, of which only 941 subjects had all four resting-state functional and diffusion MRI scans. We used these 941 in the heritability analyses in the main paper and 645 of these subjects that were white and non-Hispanic in the subgroup analysis included in Figure S5. A set of 41 of the 941 had another visit 6 months after the initial visit, which comprised the group of individuals in the test-retest analysis. An unrelated subset of 420 out of the 941 were randomly chosen for the calculation of the SC-FC coupling; 415 out of this set of 420 had composite cognition scores and were included in the GLM analysis. A second set of 346 unrelated individuals (non-overlapping with the 420) was selected for the out-of-sample validation study.

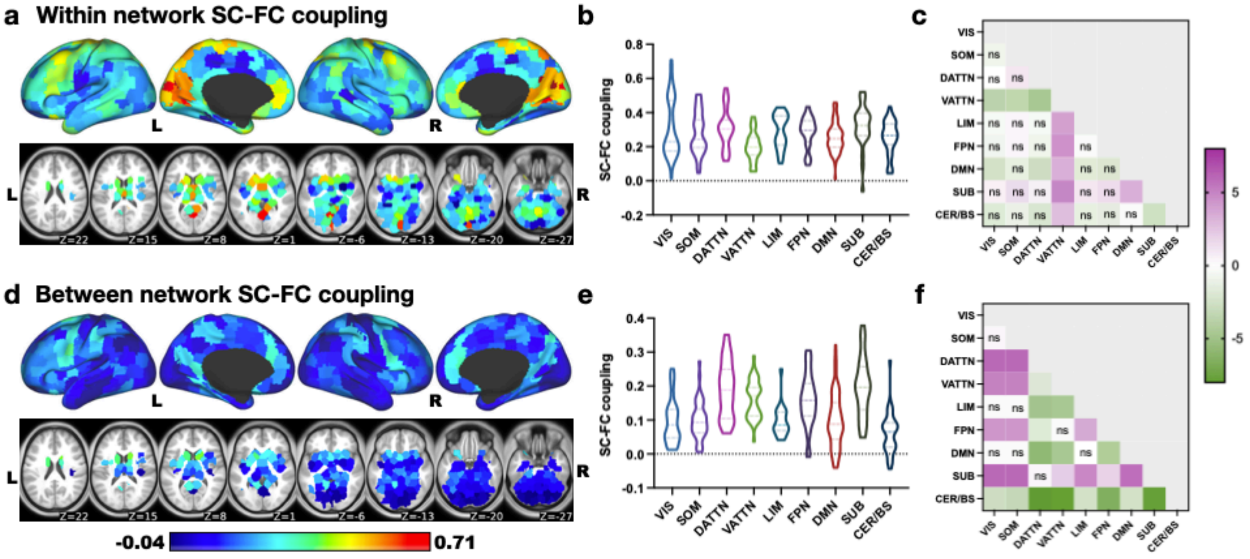


**Figure S2: Within network and between network SC-FC coupling. a** Within-network SC-FC coupling for each region is the Spearman correlation of the structural and functional connections between that region and other regions in the same network **b** Within network SC-FC coupling for the nine different networks. **c** Pair-wise comparisons of the within network coupling. **d** Between-network SC-FC coupling for each region is the Spearman correlation of the structural and functional connections between that region and other regions outside of its assigned network. **e** Between network SC-FC coupling in nine different networks. **f** Pair-wise comparisons of the between network coupling values. Dorsal/ventral attention and subcortical areas have significantly higher between-network coupling than other networks while cerebellum and brain stem have significantly lower between-network coupling than other networks.

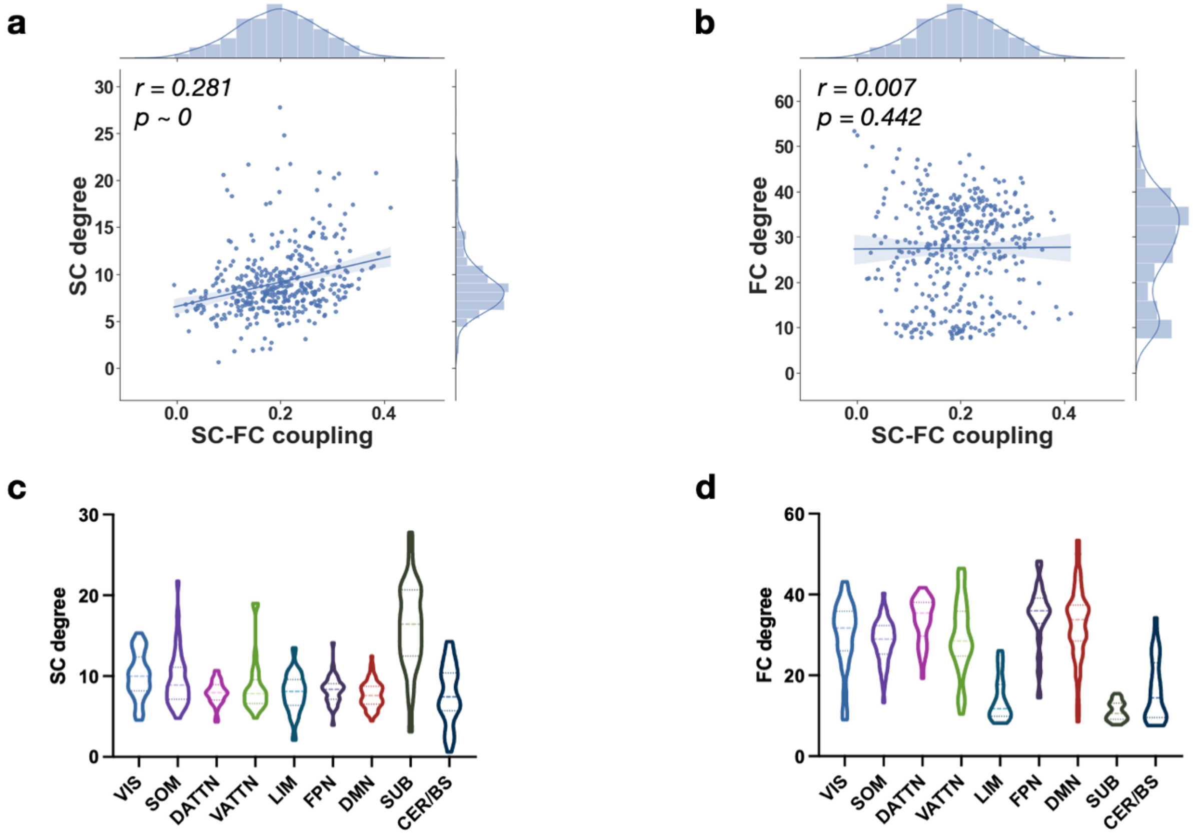


**Figure S3: Relationship of SC-FC coupling with SC and FC. a** Scatter plot of SC-FC coupling with the degree of SC. SC-FC coupling has a moderate positive correlation with the degree of SC. **b** Scatter plot of SC-FC coupling with the degree of FC. SC-FC coupling has no significant correlation with the degree of FC. **c** SC node degree by network. **d** FC node degree by network.

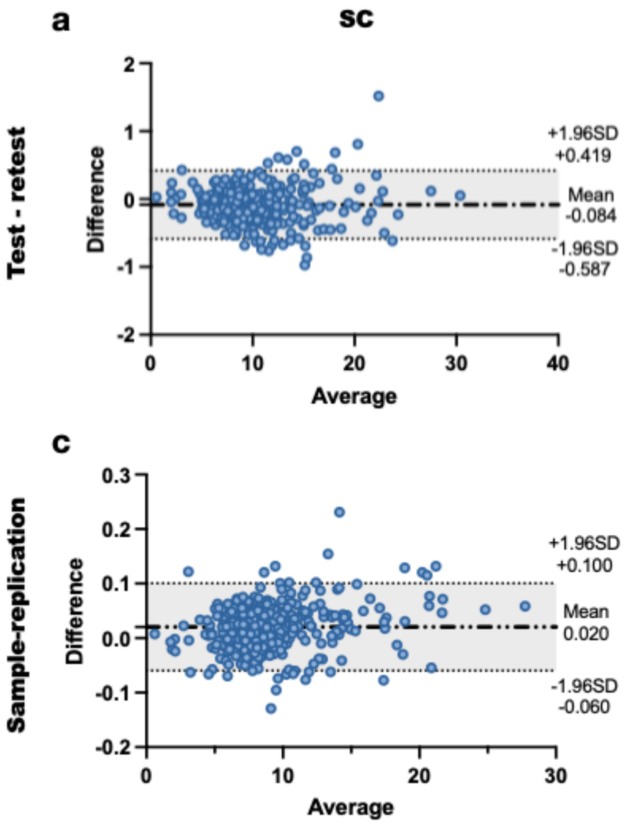


**Figure S4: Test-retest and out-of-sample reliability of SC and FC node strength.** Bland-Altmann plots of the average of the two measures (test/retest or sample/out-of-sample) against the difference in the two measures. We see generally that the two measures are quite reliable across time and across populations (test-retest Pearson correlation for FC and SC was *r=0.995* and *r=0.998*, sample and out-of-sample Pearson correlation for FC and SC was *r=0.999* and *r=0.999*).

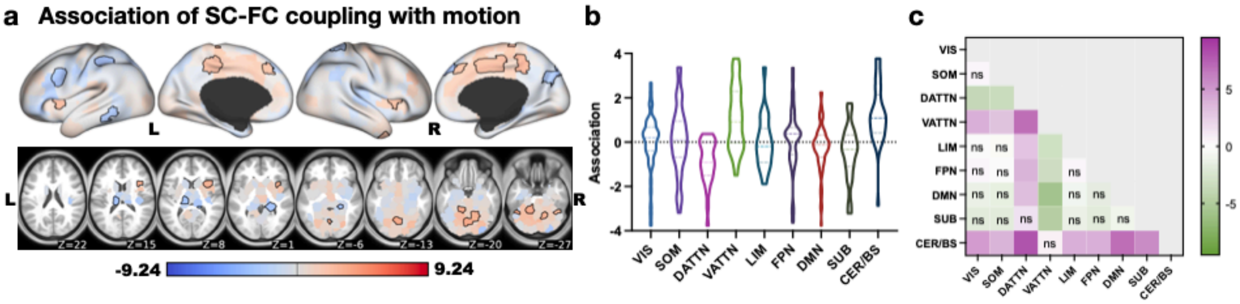


**Figure S5: Associations between SC-FC coupling and motion. a** display regional $\beta$ values from the GLM quantifying associations between SC-FC coupling and motion. Areas with significant $\beta$ values (after correction) are outlined in black. **b** show the network-wise $\beta$ values for motion. **c f i** show the t-statistics for all pairwise comparisons. Those comparisons with FDR corrected p>0.05 are marked with “ns”.

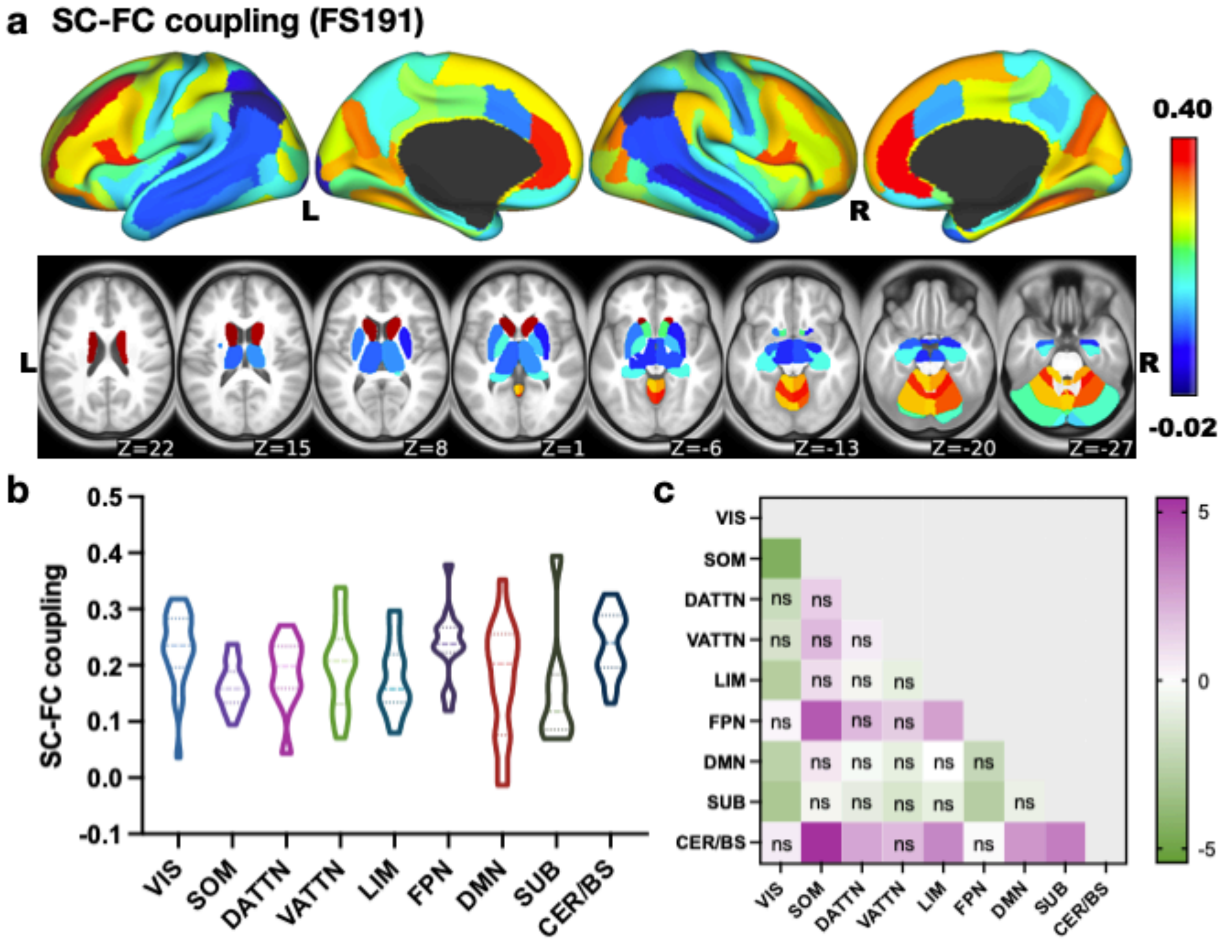


**Figure S6: SC-FC coupling in FS191 atlas. a** SC-FC coupling in FS191 atlas varies across cortical and subcortical areas with range -0.02 to 0.40. **b** SC-FC coupling distribution in nine networks. Visual, frontal parietal network, cerebellum and brain stem had generally higher coupling than other areas, with mean coupling 0.23 $\pm$ 0.06, 0.24 $\pm$ 0.06 and 0.24 $\pm$ 0.06, respectively. **c** Pair-wise comparisons between networks.

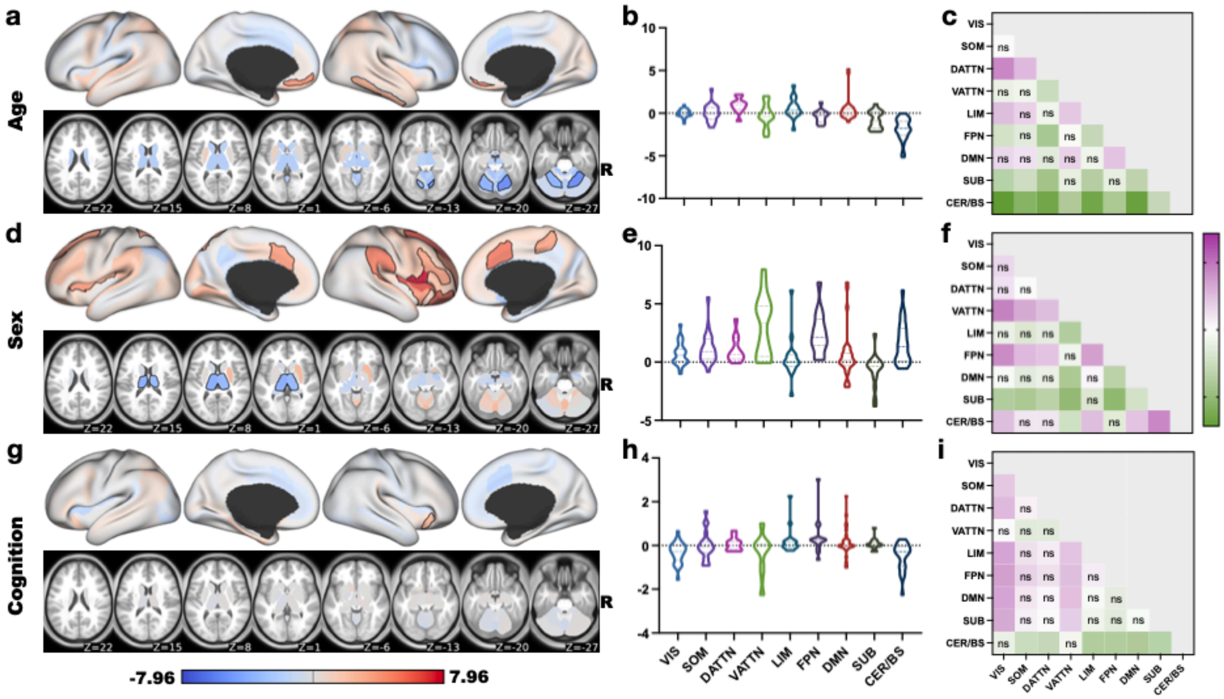


**Figure S7: Associations between SC-FC coupling (FS191) and age, sex, total cognition. a d g** display regional $\beta$ values from the GLM quantifying associations between SC-FC coupling and age, sex (blue indicates higher SC-FC coupling in females, red higher in males) and total cognition, respectively. Areas with significant $\beta$ values (after correction) are outlined in black. **b e h** show the network-wise $\beta$ values for age, sex and total cognition, respectively. **c f i** show the t-statistics for all pairwise comparisons. Those comparisons with FDR corrected p>0.05 are marked with “ns”.

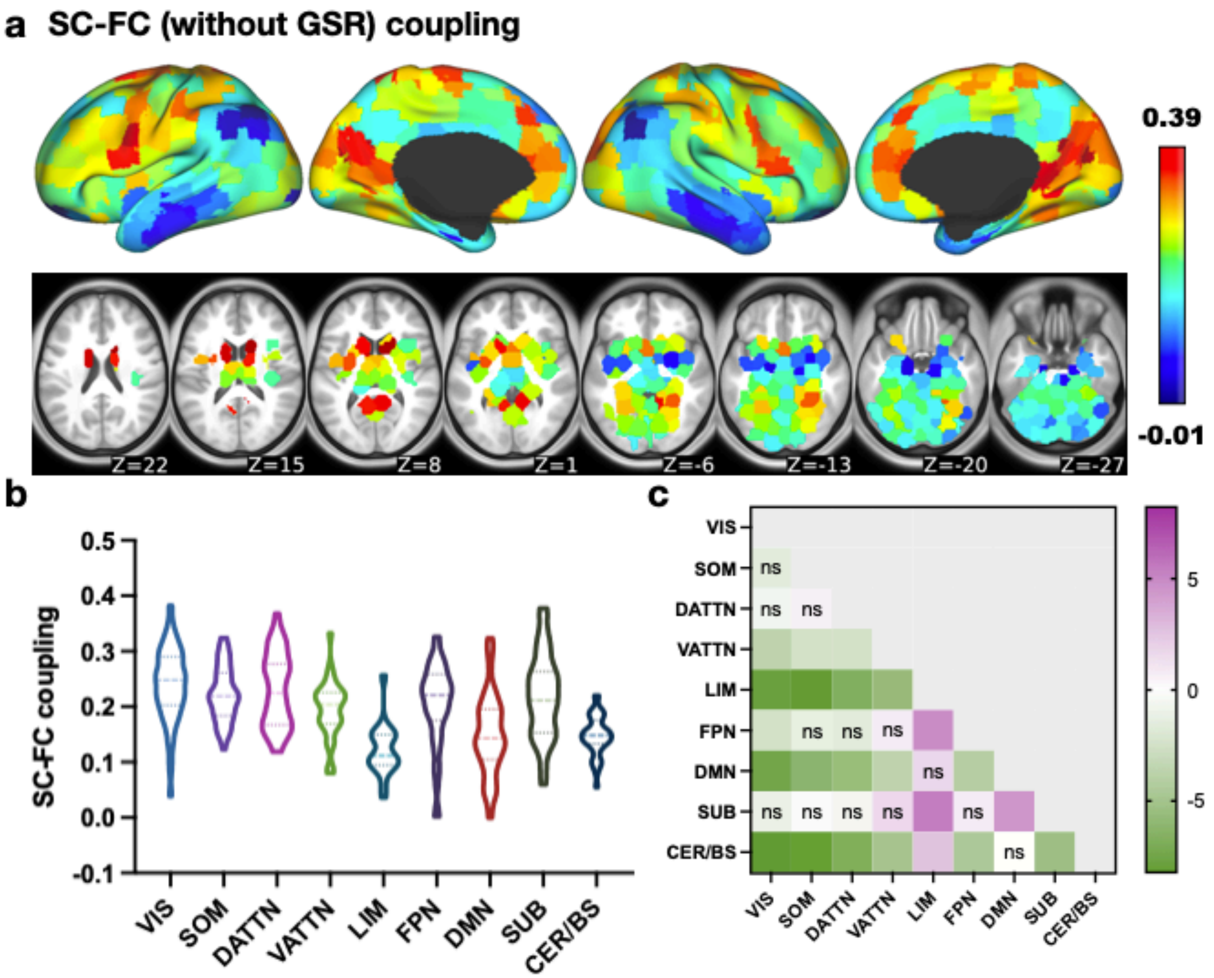


**Figure S8: SC-FC coupling computed using FC without global signal regression (GSR). a** SC-FC (without GSR) coupling varies across cortical and subcortical areas with range from -0.01 to 0.39. b SC-FC (without GSR) coupling in nine networks. **c** Pair-wise comparisons between nine networks. Pearson's correlation of the SC-FC coupling results in the main paper (with GSR) with the non-GSR coupling was *r=0.961* (*p=0*).

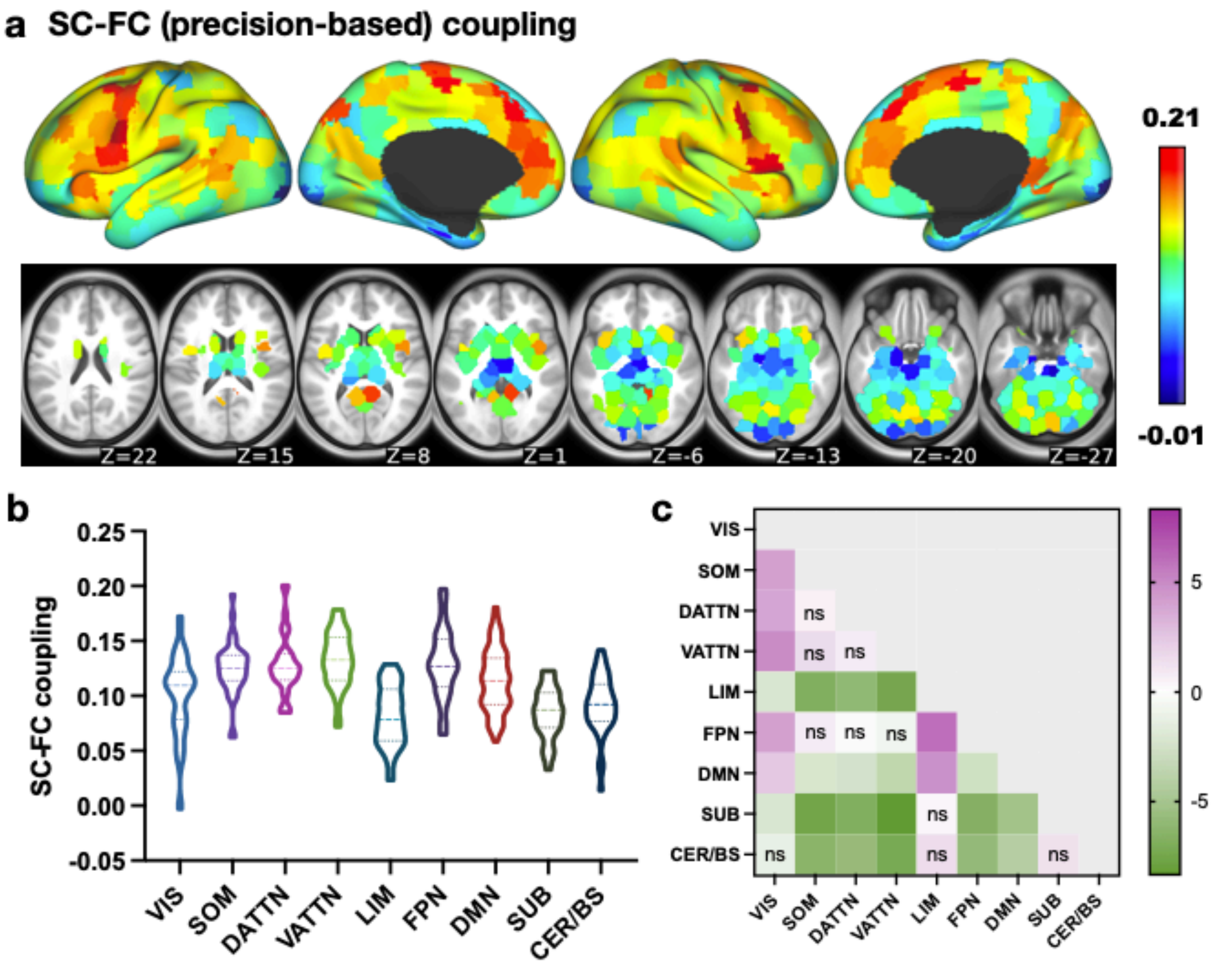


**Figure S9: SC-FC coupling calculated using precision-based FC. a** SC-FC coupling across cortical, subcortical and cerebellar regions. **b** SC-FC coupling distribution across nine networks. Ventral/dorsal attention, frontal parietal and somatomotor networks had generally higher coupling than other areas. Limbic and subcortical area had weaker mean coupling ($0.08 \pm 0.03$ and $0.08 \pm 0.02$, respectively). **c** Pair-wise comparisons between network coupling. Pearson's correlation of the SC-FC coupling results in the main paper (using full correlation-based FC) with the precision-based FC measure is *r=0.486* (*p=0*).

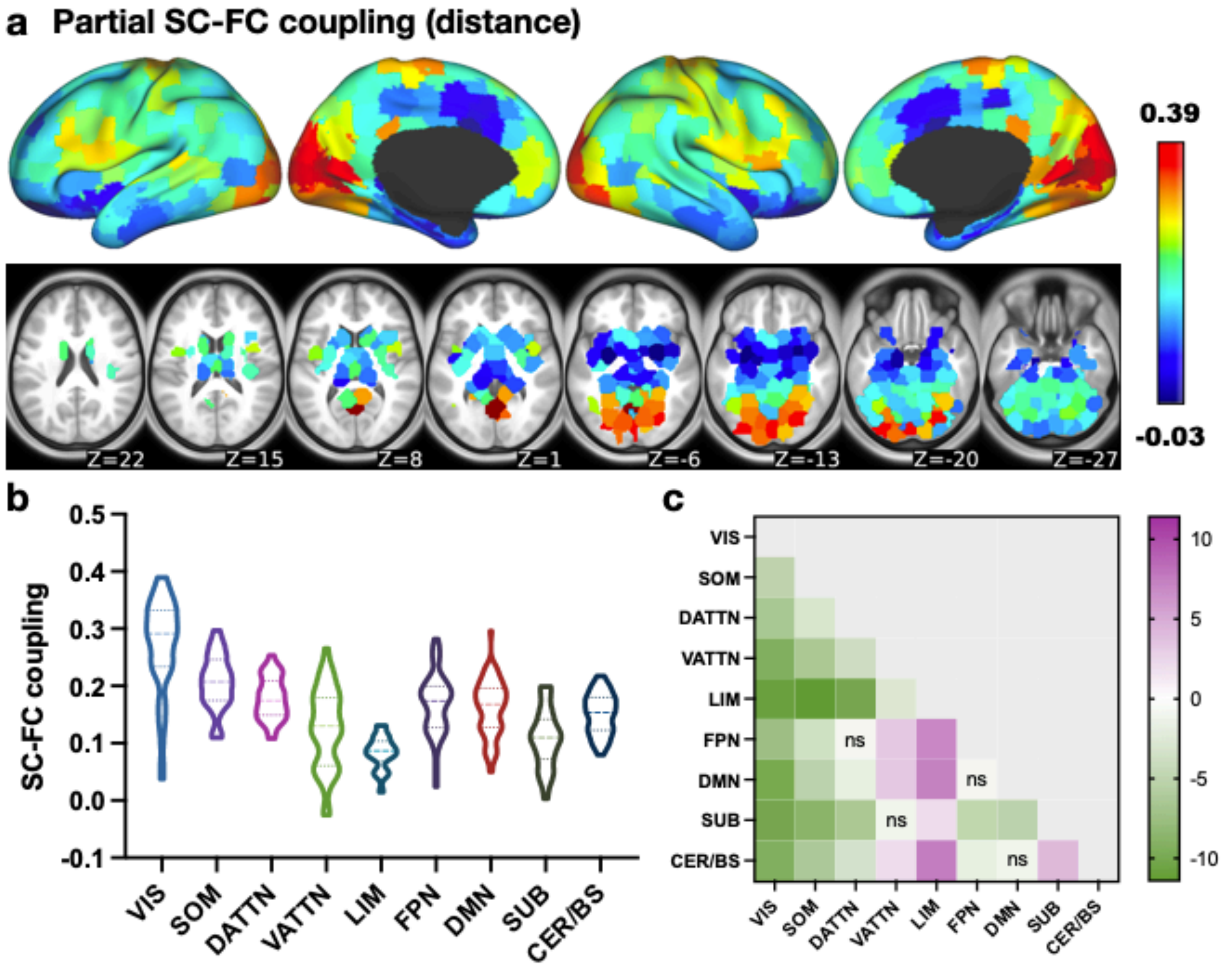


**Figure S10: Partial SC-FC coupling with inter-node Euclidean distance as a covariate. a** Partial SC-FC coupling was computed by partial Spearman correlation of the row in SC and its corresponding row in FC with the Euclidean distance between regional centroid pairs as a covariate. SC-FC coupling measured in this way varies across cortical and subcortical areas and ranges from -0.03 to 0.39. **b** Partial SC-FC coupling in nine networks. **c** Pair-wise comparisons between nine networks. Visual and somatomotor network have significant higher partial SC-FC coupling than other networks. Limbic network has significantly weaker partial SC-FC coupling. Partial SC-FC coupling is correlated with the standard SC-FC coupling (Pearson's *r=0.431, p=0*).

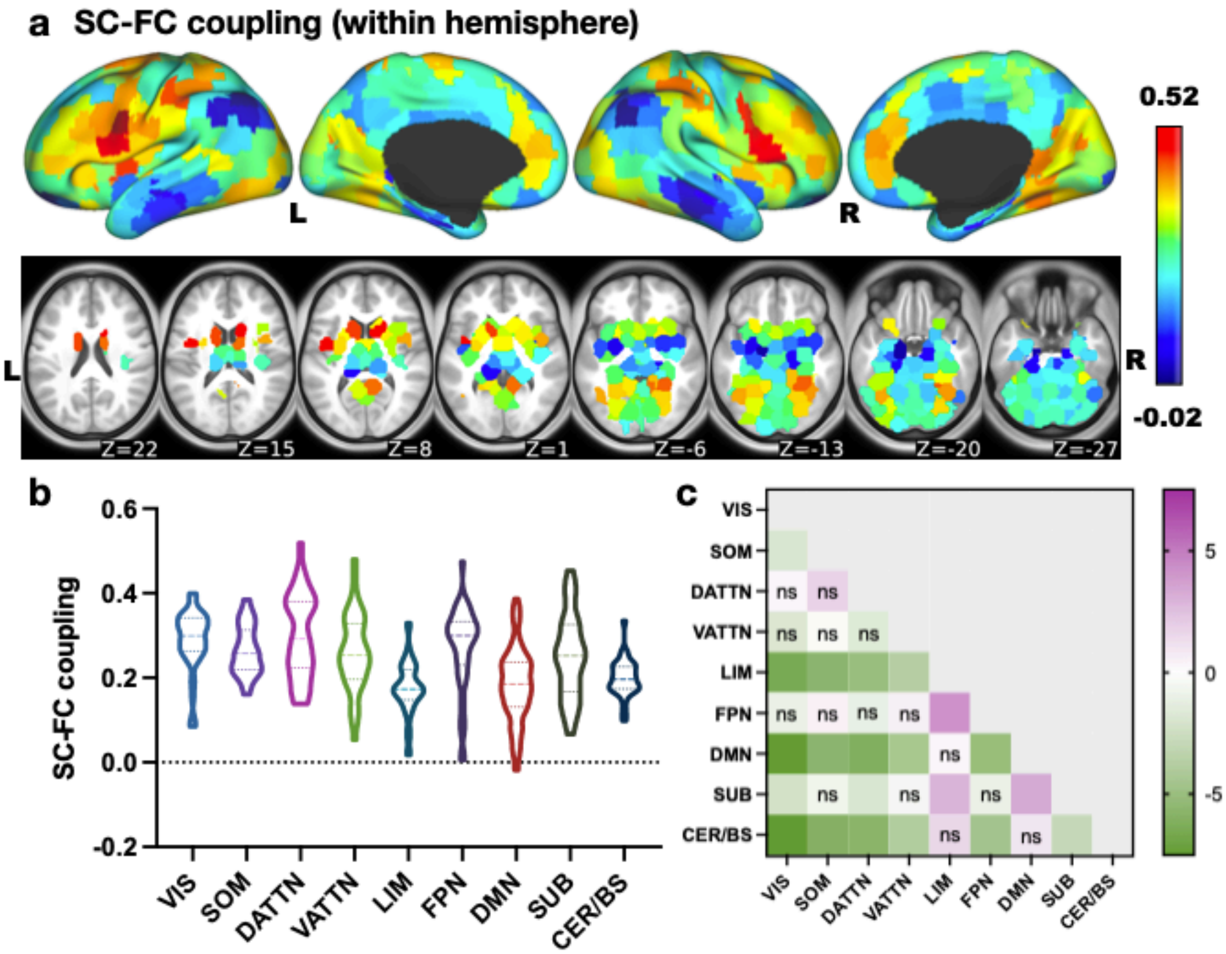


**Figure S11: SC-FC coupling within a single hemisphere. a** SC-FC within hemisphere coupling varies across cortical and subcortical areas with range from -0.02 to 0.52, which is a bit higher than whole brain SC-FC coupling but preserves consistency with the whole-brain results **b** SC-FC within hemisphere coupling in nine networks. **c** Pair-wise comparisons between nine networks. Pearson's correlation of the SC-FC coupling results in the main paper (using whole-brain SC/FC) with the single hemisphere SC/FC coupling is *r=0.864* (*p=0*).

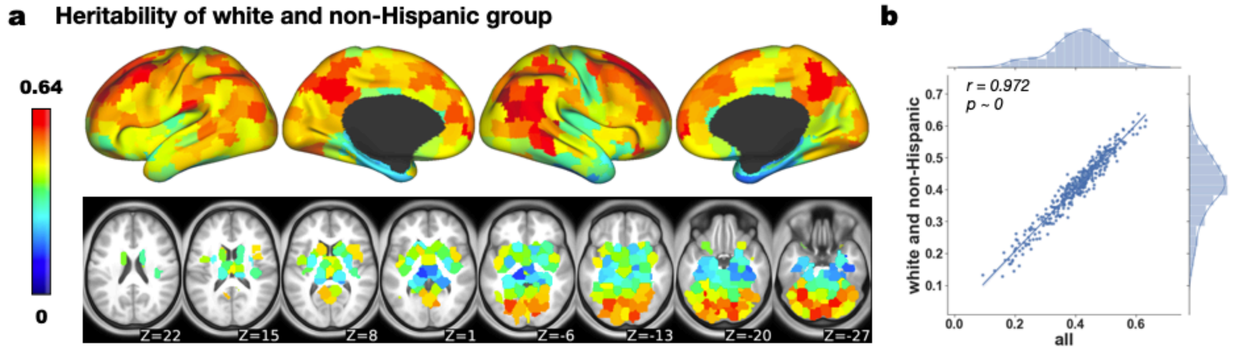


**Figure S12: Heritability of the homogenous sub-group: white and non-Hispanic group in HCP.** **a** Heritability of white and non-Hispanic group (n=645) ranges from 0 to 0.64. **b** The subgroup heritability is highly correlated (Pearson's *r=0.972, p=0*) with the heritability from all subjects presented in the main paper.
